# First evidence of nocardiosis in farmed red drum (*Sciaenops ocellatus*, Linnaeus) caused by *Nocardia seriolae* in the Gulf of Mexico

**DOI:** 10.1101/2021.01.15.426713

**Authors:** Rodolfo Enrique del Rio-Rodriguez, Jose Gustavo Ramirez-Paredes, Sonia Araceli Soto-Rodriguez, Yechiam Shapira, Mariana del Jesus Huchin-Cortes, Judith Ruiz-Hernandez, Monica Isela Gomez-Solano, David Haydon

**Author notes:** Correspondence: Sonia Araceli Soto-Rodriguez, Centro de Investigación en Alimentación y Desarrollo, Unidad Mazatlán en Acuicultura y Manejo Ambiental, Av. Sábalo-Cerritos, Mazatlán, Sinaloa, México.

## Abstract

Between August and December 2013, the offshore cages of a commercial marine farm culturing red drum *Sciaenops ocellatus* in Campeche Bay were affected by an outbreak of an ulcerative granulomatous disease, reaching 70% cumulative mortality. Thirty-one adults displaying open ulcers on the skin were submitted to our laboratory for diagnosis. Multiple white-yellowish nodules (0.1-0.5 cm in diameter) were present in all internal organs, where the kidney and the spleen were the most severely affected. Histopathological observations of these organs evinced typical granulomatous formations. Gram and Ziehl-Neelsen stains on tissue imprints, bacterial swabs and tissue sections revealed Gram-positive, acid-fast, branching beaded long rod filamentous bacteria. Tissue samples resulted positive for nocardiosis when a *Nocardia* genus specific nested PCR was used. Definite identification at the species level and taxonomic positioning of the fastidious pathogen was achieved through a specific *Nocardia seriolae* PCR and by sequencing the *gyrB* gene of pure isolates. After application of antibiotics during fry production, a posterior follow-up monitoring (from 2014 to 2017) detected mild but recurrent outbreaks of the bacteria with no seasonality pattern. To the extent of our current knowledge, this is the first report of piscine nocardiosis (in a new host -red drum) in Mexico.

## INTRODUCTION

The Food and Agriculture Organization of the United Nations describes red drum (*Sciaenops ocellatus*) as a relatively hardy, popular food fish that adapts well to captive conditions. Red drum natural distribution in the Gulf of Mexico spans from Cape Cod, in Massachusetts, USA to Tuxpan in Veracruz Mexico (FAO, 2013).

First efforts to start farming red drum began in the late 1970’s in the USA with the aims of providing alternatives to wild-caught fish, initiating stock enhancement programmes in the coast of Florida and Texas, and to recover the natural populations declining due to the heavy pressure exerted by commercial and recreational fishing activities (https://bit.ly/3sdlbAV). Since then, the farming of red drum has extended globally with China, Israel, Mauritius, Mayotte and the United States of America as the current main producers. The production of farmed red drum has not only expanded geographically but also intensively and steadily over the time passing from a discrete 198 tons in 1998 to over 71,000 tons in 2016 (FAO, 2017).

In Mexico, marine aquaculture is dominated by shrimp with 229,800 tons in 2018, that is mostly farmed in the northwestern states of the Pacific Coast, including Sonora, Sinaloa, Nayarit and Baja California Sur (Tellez-Castañeda, 2019). Finfish marine aquaculture, however, is still incipient, although several candidate species are subject of intensive research by academic and government institutions both at the Pacific and Gulf of Mexico coasts, including *Seriola lalandi*, *Seriola rivoliana*, *Centropomus viridis*, *Centropomus undecimalis*, *Morone saxatilis*, *Paralichthys californicus, Totoaba macdonaldi* amongst others (Aviles-Quevedo & Castelló-Orvay, 2004; FAO, 2013; Rueda-López et al 2017; Dávila-Camacho 2019).

In 2011, a joint venture of Israeli and Mexican entrepreneurships started a commercial farm of red drum in the state of Campeche. The farm set up located in Seybaplaya Campeche comprised an in-land “hatchery” (exact location at https://w3w.co/suben.valoren.limitantes) with semi recirculation systems and broodstock imported from the USA, a nursey section with inshore floating cages (1.5 km) for pre-growth stages and various offshore (34 km) semi-submersible cages for the grow-out phase.

Since conditions in the area were suitable for growing out all year, batches supplied by the hatchery, a smaller local producer and stock imported from Israel were often deployed mixed to the sea cages to maximize production. At the end of the production cycle (24-30 months) fish were harvested at 1-1.2 kg, eviscerated, quick-frozen and transported in ice to a local processing plant for filleting and packaging. Harvesting was escalated through the year to secure continued supply to the food chain. Processed fillets were frozen and exported to the USA. During operation, no other aquaculture facility like this existed in the Mexican side of the Gulf of Mexico.

In August 2013, the farm divers reported an outbreak of disease in the offshore cages as unusual patterns of mortalities appeared. Moribund fish displayed erratic swimming with cutaneous ulcers, loss of scales and nodular formations mainly in the dorsal area. By November 2013, the estimated cumulative mortality in the affected cages was around 70%. For the 2014 production cycles, mortalities decreased abruptly to approximately 6%, likely due to the implementation of antibiotic therapy with doxycycline but remained sustained throughout the year. For the 2015, 2016 and 2017 cycles, mortality significantly decreased with sporadic outbreaks of the disease with low mortalities. The farm finally stopped operations in early 2018, allegedly due to internal managerial problems.

In farmed marine fish, outbreaks of systemic granulomatous disease are often associated with infections of facultative intracellular bacteria (Martínez-Lara et al., 2020). The bacterial pathogens most commonly linked to granulomatous disease in fish are: *Edwardsiella* spp. (Reichley et al., 2017), *Francisella* spp. (Ramirez-Paredes et al., 2020), *Piscirickettsia salmonis* (Fryer & Lannan, 2015), *Nocardia* spp. (Woo & Bruno, 2011; Labrie et al., 2008; Cornwell et al., 2011) and *Mycobacterium* spp. (Aubry et al., 2017). The anamnesis, gross pathology and cytopathology (i.e. tissue imprints) observed during the outbreaks of disease in the red drum fish farmed in Campeche Mexico clearly warranted inclusion of nocardiosis and mycobacteriosis as a differential diagnosis. The posterior use of genus specific PCR confirmed the presence of a member of the genus *Nocardia* in the tissues of the diseased fish.

In aquaculture *Nocardia asteroides*, *N. seriolae* and *N*. *salmonicida* are the main species related with fish nocardiosis (Maekava et al., 2017). These Gram-positive, aerobic filamentous bacteria can act as opportunistic or primary pathogens in wild and farmed fish (Labrie et al., 2008). For decades, outbreaks of nocardiosis in farmed marine fish have been reported throughout East Asia i.e., China, Taiwan and Japan (Nayak & Nakanishi, 2016) and in more recent times, outside this geographic area in Australia (Bransden et al., 2000), USA (Cornwell et al., 2011), and Greece (Elkesh et al., 2013) causing considerable direct and indirect economic losses.

In the present study, the preliminary diagnosis of the initial outbreak, led to the posterior isolation, characterisation, and molecular identification of *Nocardia seriolae* as the aetiological agent. In addition to this, a series of surveillance programmes were established to monitor the progression of the disease during the following 3.5 years with the objective to control and prevent further acute outbreaks. To the best of our knowledge, this is the first clinical description of nocardiosis in aquatic organisms in Mexico. The findings of our study highlight the lack of sufficient professionals formed in fish pathology and the need of a strategy for the establishment of stringent surveillance programmes for the prevention, early detection, control and eradication of emerging diseases in the incipient finfish marine aquaculture industry in Mexico.

## MATERIAL AND METHODS

### Sample submissions and gross pathology

Between late August and early November 2013, 31 diseased grow-out fish (12-18 months old; average weight 898 g) from the offshore cages were collected alive and transported into three separate batches (n=7 on 26.08.13; n=12 on 27.09.13; n=12 on 10.12.13) to the Laboratory of Aquaculture and Aquatic Animal Health (EPOMEX Institute, University of Campeche) for diagnosis. Most specimens displayed white nodules in the gills as well as large open ulcers and/or adhesions on dorsum and flanks (Fig.1a and b). Upon arrival, fish were euthanized with a lethal overdose of tricaine methanesulfonate 1,000 mg/g, and necropsied. Samples of the affected organs were collected for histopathology, tissue imprints, molecular diagnosis and bacteriology.

**Figure 1.**
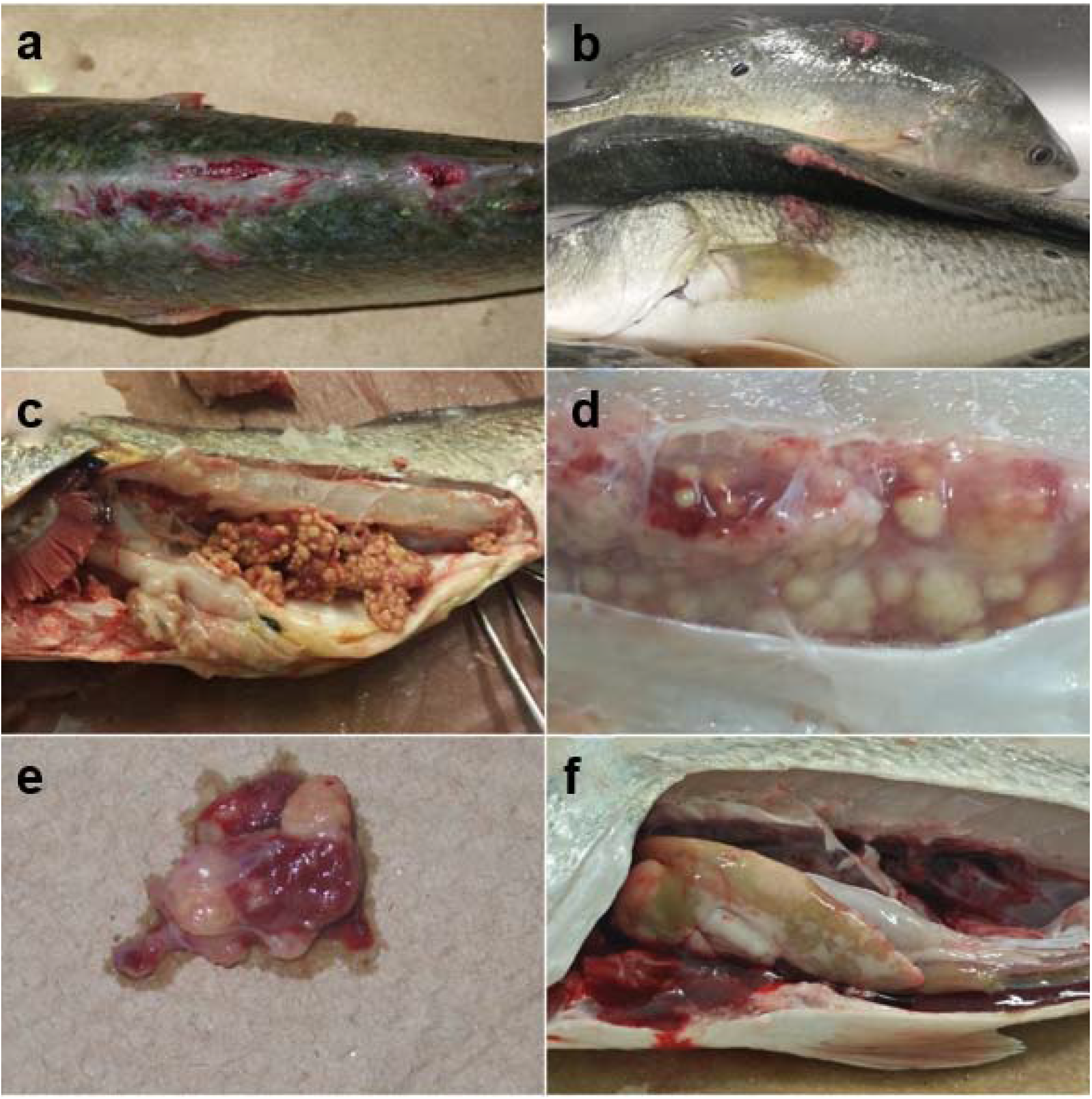
Gross pathology observed in diseased red drum farmed in Campeche Mexico during outbreaks of nocardiosis. (**a**) Dorsal view red drum displaying open ulcers due to *Nocardia seriolae*. (**b**) Young adult red drum specimens with flank and dorsal granulomatous lesions. (**c**) A severe case of granulomatous formations in kidney in a young red drum adult specimen; kidney was dissected off to display granulomas. (**d**) Close up of nodular formations seen in (c) before dissection of the kidney. (**e**) Dissected heart of a diseased red drum with coalescent granulomas and polyserositis formation. (**f**) Fibromatous congestion and lipoidosis of the liver (centre of the picture) of a specimen positive for *Nocardia seriolae*.

### Cytopathology (tissue imprints)

Organ imprints were made onto clean microscope glass slides from cut-through sections of affected kidney, spleen, and heart. The impression prints were air dried for 1-2 minutes and then stained with Gram, Ziehl–Neelsen (ZN) and Fite-Faraco stains following the kit’s manufacturer instructions.

### Histopathology

Samples of muscle of approximately 1 cm^3^ with ulcerated skin, kidney, heart, spleen and liver with nodular lesions were fixed in 10% (v/v) buffered formalin during 48h prior to processing using standard protocols for histology, embedded in paraffin wax and sectioned at 5 μm on a Leica™ microtome. Slides were left overnight in an oven at 40°C to ensure adhesion. Thereafter, sections were stained with haematoxylin and eosin (H&E); some sections were stained with ZN and Fite-Faraco stains and examined under a light microscope. Digital images were obtained using a Leica Digital Camera MC 120HD and capture software.

### Molecular diagnosis in fish tissues

To conduct the molecular diagnosis, samples of 25 mg of tissue were fixed, and 15-30 μg of DNA was extracted from ethanol-fixed fish tissues using a DNeasy Blood & Tissue Kit (Qiagen) following the manufacturer’s guidelines.

For mycobacteriosis extracted DNA was used a template on a *Mycobacterium* genus specific PCR using the primers T39 5’-CGAACGGGTGAGTAACACG-3’ and T13 5’-TGCACACAGGCCACAAGGGA-3’ as per the protocol by Talaat et al. (1997).

For nocardiosis extracted DNA was used as template on a *Nocardia* genus specific nested PCR as per the protocol by Elkesh *et al.* (2013). For the first PCR, universal 16S rRNA primers i.e. 20F (5’-AGAGTTGATCATGGCTCAG-3’) and 1500R (5’-CGATCCTACTTGCGTAG-3’) (Weisberg et al. 1991) were used. The PCR reaction consisted of 12.5 μL HotStarTaq® Master Mix (Qiagen) [200 μL of each dNTP, 1.5 mM MgCl_2_], 0.5 μL of each primer, 6.5 μL RNase free water and 5 μL of DNA template (final volume of 25 μL). The amplification reaction was performed in a DNA thermocycler (Applied Biosystems®) with minor modifications in the cycling conditions. Briefly, initial denaturation for 15 min at 95°C was followed by 35 cycles of denaturation at 94°C for 1 min, annealing at 62°C for 1 min, extension at 72°C for 1 min with a final extension at 72°C for 10 min. For the second PCR, 5 μL of the initial PCR product per sample were used as a template with *Nocardia* genus specific primers i.e., NG1 (5’ACCGACCACAGGGGG-3’) and NG2 (5’-GGTTGTAAACCTCTTTCGA-3’) and HotStarTaq® Master Mix Kit (Qiagen) up to a final volume of 25 μL. For amplification, the following minor modifications in the cycling conditions were adopted: initial denaturing for 15 min at 95°C followed by 30 cycles of 94°C for 1 min, 62°C for 1 min and 72°C for 1 min with a final extension of 72°C for 10 min. The amplification products of the second PCR were visualized after electrophoresis (35 min at 90 V) of 5μL per sample on 2.0% agarose gel stained with ethidium bromide using Tris-acetate-EDTA buffer (40 mM Tris, 20 mM acetic acid, and 1 mM EDTA, pH 8.0) and a 100-bp GelPilot® DNA Molecular Weight Marker (Qiagen).

### Bacterial isolation and phenotypic characterisation

Loop swabs were taken from head or trunk kidney and/or spleen where nodules were more evident and streaked onto tryptic soy agar (TSA), TSA + 1.5% NaCl, and slopes of Lowesten-Jensen medium (LJM). Plates and slants were incubated at 27°C for 15 days with daily observations and colonies were examined by Gram and ZN staining. Acid-fast positive rods were tested for catalase and oxidase production and subcultured on TSA and brain heart infusion (BHI) agar at 0, 1, 1.5, 2, 3, 4 and 6% NaCl. Strains were also subcultured in LJM slopes with glycerol, some slopes with bacterial growth were deep-freeze (−80°C) for future testing.

### Bacterial identification by species-specific 16SrRNA PCR

The identity of one of the isolates (J8P8) recovered during the follow up visits (see “Disease Progression” section) was confirmed using a *Nocardia seriolae* species-specific PCR targeting the 16S rRNA gene, with primers NS1 5’-ACTCACAGCTCAACTGTGG-3’ and NG1: 5’-ACCGACCACAAGGGGG-3’ as per the protocol by Miyoshi & Suzuki (2003). For this, the isolate was cultured on LJM at 28 °C for 4 days. DNA extraction from bacterial colonies was performed using a Wizard® Genomic DNA Purification Kit (Promega) according to the manufacturer’s instructions. The PCR reaction was carried out in a 14.51 μL reaction mixture containing PCR buffer 1X with (NH_4_)S0_4_, 2.76 mM MgCl_2_, 0.45 m dNTP mixture, 0.45μM of each primer, 0.02 U/μL Taq polymerase (Thermo Scientific™), 50 ng/μL DNA and 1X water for qPCR. After denaturation at 94℃for 3 min (DNA thermocycler Applied Biosystems®), the amplification reaction was run through 30 cycles of denaturation at 94℃ for 1 min, annealing at 58℃ for 20 s and extension at 72℃ for 1 min. Final extension was performed at 72℃ for 10 min. A strain of *N. seriolae* recovered from diseased-farmed pompano *Trachinotus blochii* in South East Asia (Ramirez-Paredes et al unpublished data) was used as positive control. From the reaction 5 μL of the PCR product were run on 1.0 % agarose gel at 120 volts for 30 min at room temperature, using Tris-acetate-EDTA (40 mM Tris, 20 mM acetic acid, and 1 mM EDTA, pH 8.0) buffer using a 100-bp GelPilot® DNA Molecular Weight Marker (Qiagen) and visualized by Axygen® Gel Documentation System.

### Bacterial identification by *gyrB* gene sequencing

Bacterial species identification was performed on two isolates i.e. TW294/20 and TW295/20 derived from the representative strain J8P8 by targeting the subunit B protein of DNA gyrase (topoisomerase type II), i.e. the *gyrB* gene (Yamamoto & Harayama, 1995). Total nucleic acids were extracted from a loopful of pure colonies using nanomagnetic beads i.e. Genesig Easy DNA/RNA Extraction Kit (Primerdesign, Southampton, UK). PCR reactions consisted of each primer at 10 μM, 1 unit of GoTaqG2 master mix (Promega UK), 5 μL of DNA sample and milliQ water to reach a final reaction volume of 25 μL. The following thermal cycling conditions were used in a G-storm thermocycler: initial denaturing of 95°C for 5 min; 35 cycles of denaturing at 95°C for 30 s, annealing at 55°C for 30 s and extension of 73°C for 1 min; followed by a final elongation step at 73°C for 7 min. The PCR products were visualized on a 1% agarose gel stained with ethidium bromide, then purified using a QIAquick PCR Purification Kit (Qiagen) as described by the manufacturer. Then 3.5 μL of the clean-up were mixed with 2.5 μL of each of the forward and reverse primers in a separate nuclease free Eppendorf tube with 1.5 μL of nuclease free water to reach a total volume of 7.5 μL and sent for Sanger sequencing to GATC (Eurofins Genomics, Cologne, Germany).

The quality of the resulting chromatograms was visually checked using BioEdit® software version 7.1.11 and forward and reverse sequences assembled using the Multiple Sequence Comparison by Log-Expectation (MUSCLE) application of the MEGA (Molecular Evolutionary Genetic Analyses) package version X (Kumar et al., 2018). Consensus sequences were deposited in GenBank® and compared to known sequences in this database using the *in silico* nucleotide alignment tool ‘BLASTn®. Relevant sequences within the family *Nocardiaceae* and close related family *Mycobacteriaceae* were retrieved from GenBank® using pairwise comparisons were performed with all the sequences within and between the taxa using BLASTn® programmes.

All the relevant sequences were also aligned using the MUSCLE application of the MEGA software version X (Kumar et al., 2018). The alignment was manually adjusted and trimmed and its suitability for phylogenetic analyses double checked by computing the pairwise and the overall mean distances in MEGA package version X (Kumar et al., 2018), the NCBI accession number was indicated for all the individual sequences. The alignment was used to build phylogenetic trees and to analyse the evolutionary relationship of the novel *N. seriolae* Mexican isolates with its closest members in the genus and close related genera and families.

The evolutionary analyses were constructed in MEGA software version X (Kumar et al., 2018) using the Maximum Likelihood (ML) approach with exclusion of gaps and missing data. The model for each tree was chosen based on the best combination of model and rates among sites, and such combination was investigated for each alignment using the default settings of the “find best DNA/protein model” option. The reliability (reproducibility) of the trees was tested using the bootstrap method with 1,000 replications. In the analysis, the nearest neighbour interchange was chosen as the ML heuristic method.

### Antimicrobial sensitivity testing

The susceptibility to 7 antimicrobial compounds of three representative strains i.e. CV7Y, CV9Y and CV11Y isolated during the main outbreak (2013) was investigated using the qualitative Kirby-Bauer disk diffusion method as described by (Hudzicki, 2009). Briefly, isolates were suspended into sterile buffered saline solution (0.25% NaCl) from which serial dilutions were performed. About 200 μL of suspensions matching the density of McFarland 0.5 were streaked onto Mueller-Hinton and TSA (2% NaCl) media. Bacteria were tested to antibiotic discs that contained doxycycline 30 μg, oxytetracycline 30 μg, chloramphenicol 30 μg, florfenicol 30 μg, enrofloxacin 30 μg, sulphamethoxazole/trimethoprim 25 μg and clindamycin 2 μg. Plates were incubated at 27°C and checked at 24, 48 and 72 hours post-inoculation.

### Disease progression

A follow-up sampling scheme was agreed with the farm owners where monthly random samples (n=60) were collected and analysed from 2014 to mid-2017. All specimens collected during that period were subjected to diagnostic work, from routine inspection, bacteriology and histology to PCR at genus level (Elkesh et al. 2013).

## RESULTS

### Clinical signs, gross pathology and bacterial isolates

External signs of diseased red drum included lateral or bilateral exophthalmia, abdominal distension and desquamation, wasting and variable number of skin ulcerative lesions with adhesions in the flanks and dorsal region of the body. In addition, extremely pale gills were observed in some fish. At necropsy, numerous white-yellowish macroscopic nodular lesions either individual or coalescent from 0.1 to 0.8 cm in diameter, which were readily visible were widespread in most of the internal organs with evident extension in the kidney, heart and spleen (Fig. 1c, d, and e), while the liver was rarely affected. Sometimes nodules were observed in the peritoneum. The liver was not always compromised, although on most occasion nodules were concomitant with lipoidosis (Fig.1f). Severe cases of nodular formations in the kidney were seen, although those fish were still alive when collected. No other conspicuous pathogens were observed or isolated during the study. Occasionally, amphipod crustacean parasites of the family *Flabelligeridae* were found in the oral cavity of some specimens.

### Histopathology and Cytopathology

At tissue level, most of the lesions observed were of classical granuloma formations, although the organs affected displayed varying patterns of granuloma development. The most severe affected organ was the kidney (Fig. 2a), followed by the heart and spleen (Fig. 2b). In the worst cases the kidney displayed total loss of architecture with parenchymal tissue replaced by inflammatory and scaring-fibrotic cells forming the walls that contained individual and coalescing irregular granulomata.

**Figure 2.**
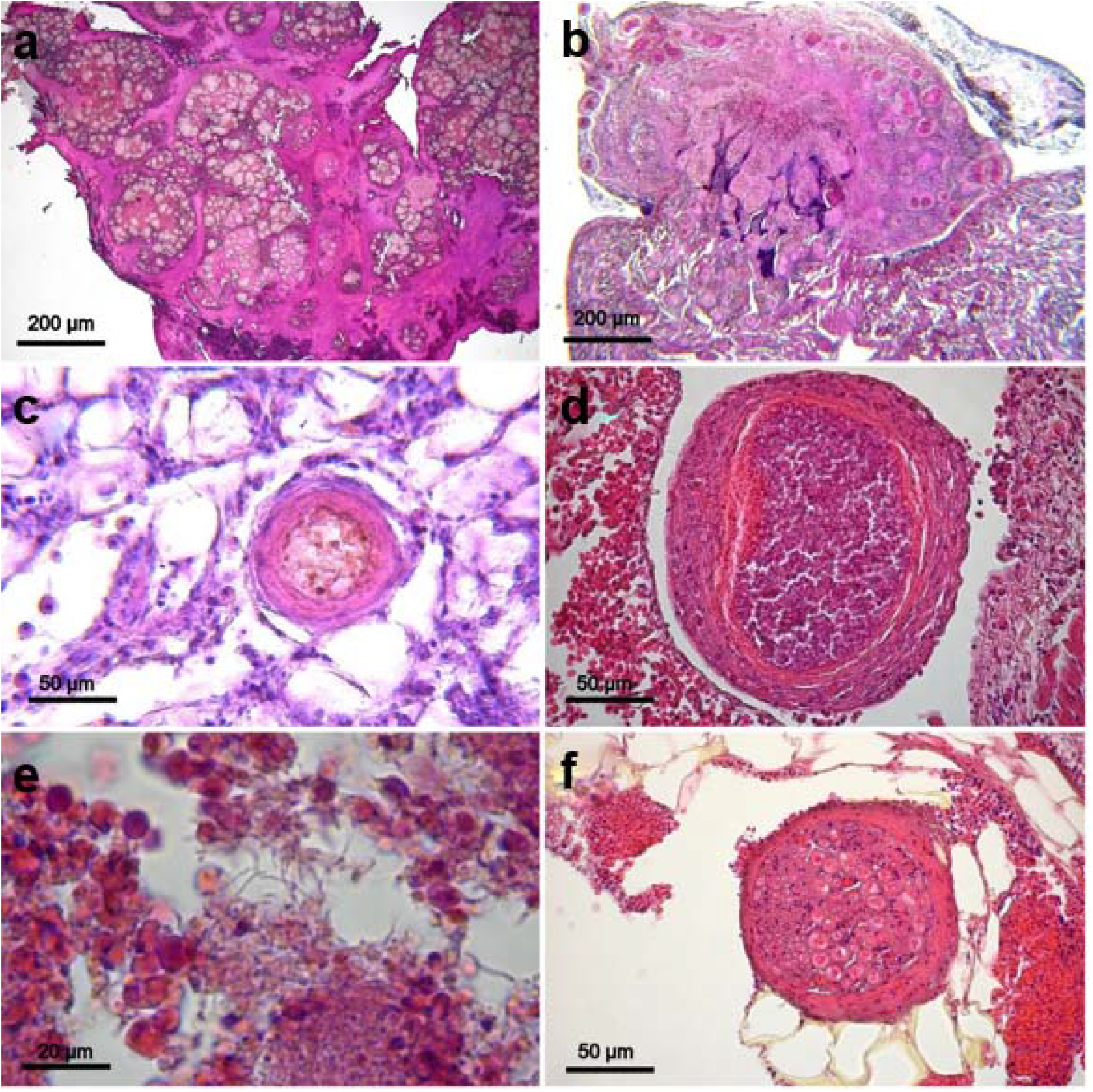
Histological sections of kidney, heart, liver and mesentery of red drum farmed in Campeche Mexico affected by piscine nocardiosis. (**a**) Section of posterior kidney with severe granulomatosis displaying total loss of architecture where parenchyma has been replaced by scaring fibrotic tissue surrounding coalescent granulomata (H&E, Light microscopy 100X). (**b**) Granulomata outgrowth from myocardial wall muscle fibres. The presence of inflammatory cells within the granuloma is extensive (H&E, Light microscopy 100X). (**c**) Small granuloma embedded in fatty storing hepatocytes. Lipoidosis was common in the cultured red drum (H&E, Light microscopy 400X). (**d**) Large granuloma between the atrium and ventricle enclosing inflammatory elements with necrotic outer cells in its vicinity (H&E, Light microscopy 400X). (**e**) Colonies of *Nocardia seriolae* among inflammatory cells. Some individual large bacterial rods can be appreciated (H&E, Light microscopy 1000X). (**f**) Granuloma in mesenteries and fat tissue with a primary core containing small secondary cores (H&E, Light microscopy 400X).

Liver sections (Fig. 2c) revealed that most of the specimens had lipoidosis, but contrary to expected, few displayed granulomas. The central zone of most granulomas had necrotic cells and debris surrounded by an inner compressed spindle-cell layer and an outer epithelioid cell layer (Fig. 2d), sometimes with visible aggregates of large slender ZN or Gram-positive rods (Fig. 2e). No giant cells were observed. Rarely, some regular granulomas found in the mesenteries had a primary core containing several small secondary cores (Fig. 2f). Granulomas of the hypodermis with growth outs to the skin were causing diverse degrees of myositis at the junction with the skin. Tissue imprints revealed the long rod body of the bacterium and branching colony morphology (Fig. 4a, b and c) of the acid-fast, Gram+ rod with beaded appearance also observable in tissues (Fig. 4e).

### Molecular diagnosis in fish tissues

No amplicon was produced using the *Mycobacterium* genus specific PCR. The *Nocardia* genus specific PCR yielded a product of ~600 bp (Fig. 3) in all of the samples tested, leading to the posterior isolation and characterization of the etiological agent.

**Figure 3.**
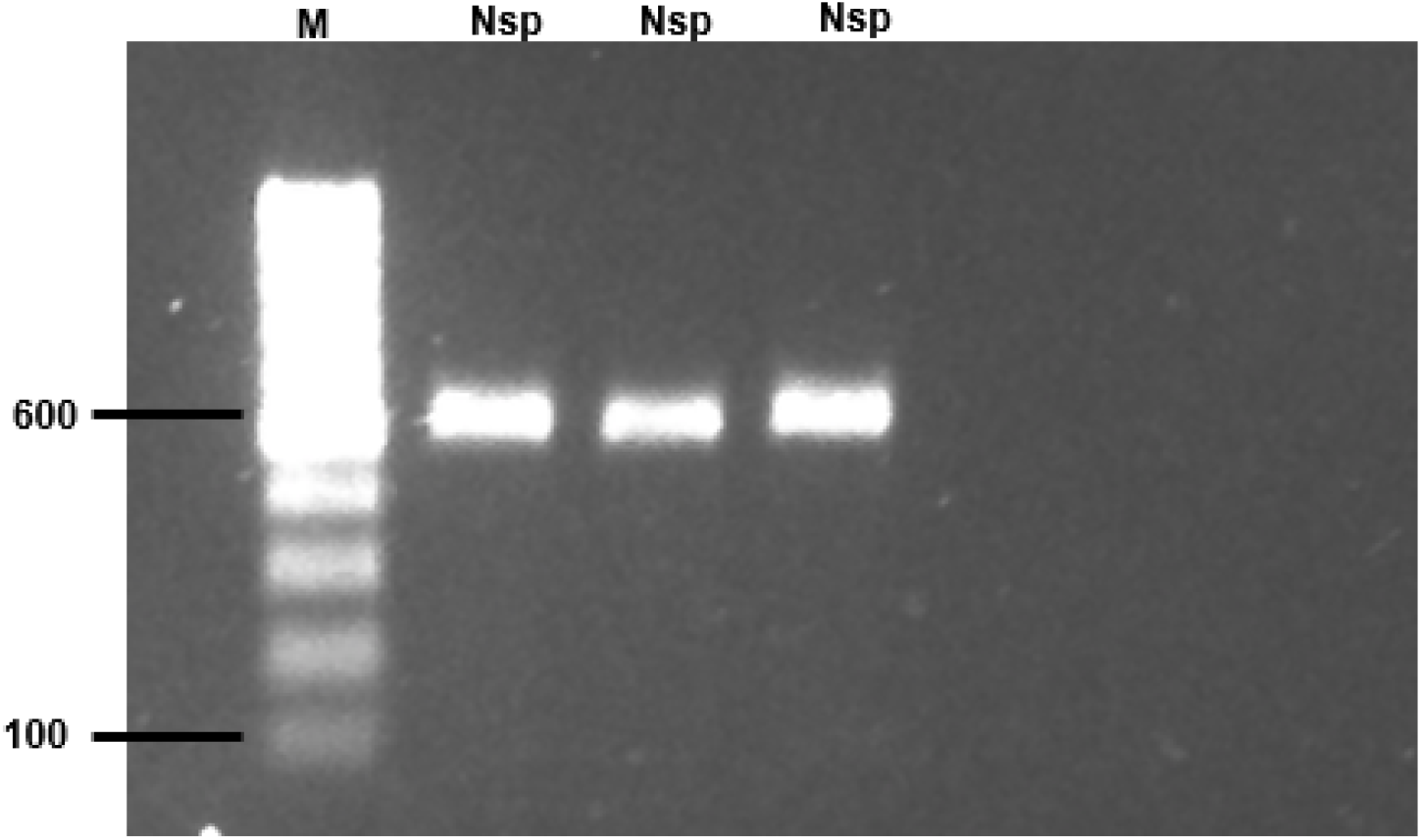
Ethidium bromide stained 1% agarose gel after electrophoresis. Molecular confirmation by PCR at genus level of ***Nocardia sp.*** from bacteria recovered form diseased fish. **M**= 100bp molecular weight markers; **Nsp**= strain J8P8 of *Nocardia sp.* by triplicate.

**Figure 4.**
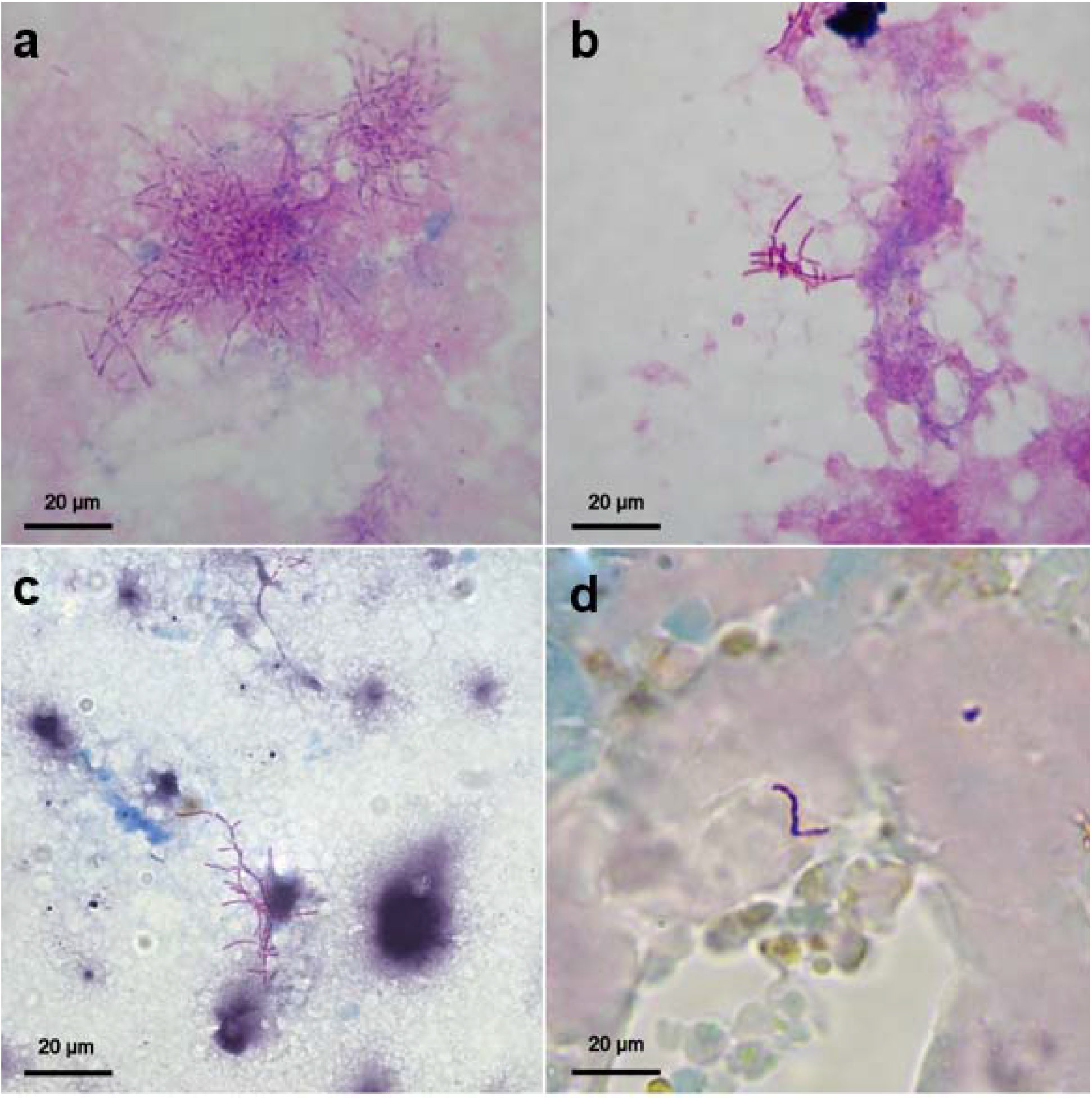
Individual and colony traits of *Nocardia seriolae* under different cell stains and histology (Light microscopy 1000X oil). (**a**) Gram+ stain of a bacterial colony of *Nocardia seriolae*. (**b**) Acid fast positive *N. seriolae* (Ziehl–Neelsen) swab from heart tissue. (**c**) Kidney swab stained with Fite-Faraco displaying branching acid-fast positive *N. seriolae*. (**d**) Histological section with Ziehl–Neelsen; partial acid-fast staining –cell wall displaying clear areas- is considered a characteristic of *Nocardia* genus that differentiates from mycobacteria. Light microscopy 1000X

### Bacterial isolation and phenotypic characterisation

Bacterial colonies were light brown in colour and with an irregular, wrinkled and sandy appearance on TSA and LJ. While being subcultured they were particularly hard and rather difficult to detach and manipulate once off from the solid media. Under the light microscope they were Gram-positive forming an irregular branching structure. Fresh tissue swabs (hearth and kidney) were also positive for acid-fast bacilli when stained with ZN and Fite-Faraco stains (Fig. 4).

Long, slender, multiseptated rods Gram and ZN positive were recovered from 14 out of 31 fish on TSA and LJM after 3 to 4 d of incubation at 28°C. Growth was observed in TSA supplemented with 1, 1.5, 2, 3 and 4% NaCl but negative with 6%. Subculture colonies growing on slants of TSA and Lowesten-Jensen media (LJM) were light brown, irregular, dry and wrinkled; senescent colonies on LJM turned chalky although some subcultures out of these colonies produced fresh growths. ZN stain of tissue imprints or colony swabs displayed individual or filamentous-branching rods with beaded appearance. Purification of some of the strains isolated in LJM was sometimes difficult as they were easily superseded by another faster growing bacterium. Some methods for separating and recovering back those strains as pure culture were not successful, namely growing on specific or general solid or broth media, media with antibiotics and antifungal compounds or incubation at higher temperatures (>30°C). Growths were readily obtained from samples of infected tissues cultured at 24-26°C after three to four d, although some subcultures of those growths did not produce colonies under the same temperatures after 15 d or later. However, more conspicuous growth was observed in LJM when subcultures were placed or transferred at 30°C. Although *Nocardia* sp. was initially isolated in TSA, purification or subculturing of the isolates into that media was not always successful.

### Bacterial identification by species-specific 16SrRNA PCR

The primers used to confirm the isolates as *N. seriolae* strains were successful as an amplicon of ~400 bp was observed in the gel electrophoresis (Fig. 5)

**Figure 5.**
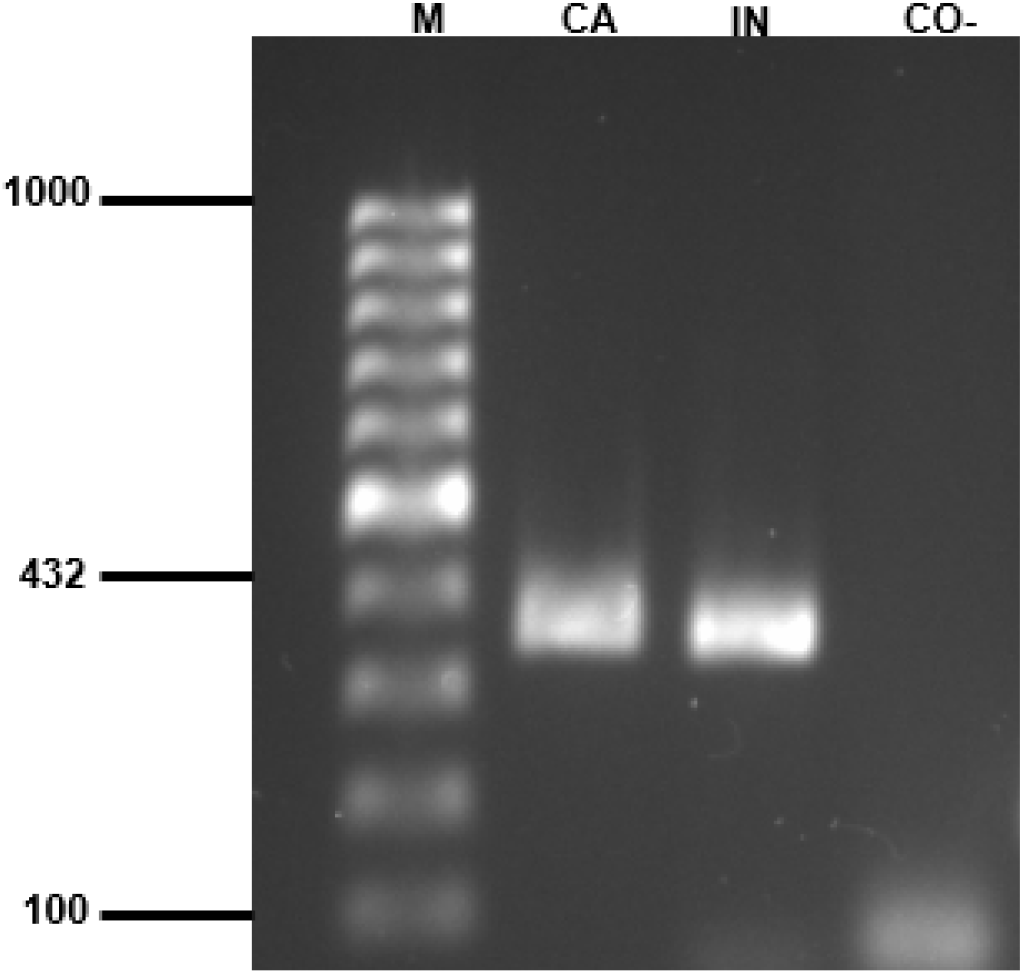
Molecular confirmation by PCR of *Nocardia seriolae*. **M**= 100bp molecular weight markers. **CA**= sample of *N. seriolae* isolated in the present study; **IN**=positive control strain of *N. seriolae* recovered from diseased-farmed pompano *Trachinotus blochii* in South East Asia (Ramirez-Paredes et al., unpublished data); **CO**-= Negative control.

### Bacterial identification by *gyrB* gene sequencing

When comparing the *gyrB* gene sequences of the isolates TW925/20 (GenBank® MW287625) and TW924/20 (GenBank® accession number # MW433931) the average nucleotide identity (ANI) between both was 100% and 99.6% similar to other *Nocardia seriolae* strains. After this, the closest related sequences were those of *Nocardia concava* (95%) and *Nocardia nigatensis* (92%), followed by *Nocardia crassostrae* (91%) and *Nocardia otitidiscaviarum* (90%). The two other known fish pathogens i.e. *Nocardia salmonicida* and *Nocardia asteriode*s were below the 90% threshold with values of 89% and 87% respectively. The similarity with *Nocardia mexicana*, a species recovered from pus samples of human mycetomas in Mexico City was also 97%. The threshold with close related genus *Skermania piniformis* was 83% and with closely related family *Mycobacteriaceae* was 80%. The complete set of pairwise comparisons is presented in Table 1.

**Table 1.** Pairwise percent (%) similarity comparisons of the *gyrB* gene.

| Bacterial species and strains | 1 | 2 | 3 | 4 | 5 | 6 | 7 | 8 | 9 | 10 | 11 | 12 | 13 | 14 | 15 | 16 | 17 | 18 | 19 | 20 | 21 | 22 | 23 | 24 |
| --- | --- | --- | --- | --- | --- | --- | --- | --- | --- | --- | --- | --- | --- | --- | --- | --- | --- | --- | --- | --- | --- | --- | --- | --- |
| [1] *Nocardia* sp. strain **TW294-20** PRESENT STUDY | - |  |  |  |  |  |  |  |  |  |  |  |  |  |  |  |  |  |  |  |  |  |  |  |
| [2] *Nocardia* sp. strain **TW295-20** PRESENT STUDY | 100 | - |  |  |  |  |  |  |  |  |  |  |  |  |  |  |  |  |  |  |  |  |  |  |
| [3] *Nocardia seriolae* strain MH196537 (CP059737.1) | 100 | 100 | - |  |  |  |  |  |  |  |  |  |  |  |  |  |  |  |  |  |  |  |  |  |
| [4] *Nocardia seriolae* strain NK201610020 (CP063662.1) | 100 | 100 | 100 | - |  |  |  |  |  |  |  |  |  |  |  |  |  |  |  |  |  |  |  |  |
| [5] *Nocardia seriolae* strain EM150506 (CP017839.1) | 100 | 100 | 100 | 100 | - |  |  |  |  |  |  |  |  |  |  |  |  |  |  |  |  |  |  |  |
| [6] *Nocardia seriolae* strain UTF1 (AP017900.1) | 100 | 100 | 100 | 100 | 100 | - |  |  |  |  |  |  |  |  |  |  |  |  |  |  |  |  |  |  |
| [7] *Nocardia seriolae* strain SB0731 (KY368575.1) | 100 | 100 | 100 | 100 | 100 | 100 | - |  |  |  |  |  |  |  |  |  |  |  |  |  |  |  |  |  |
| [8] *Nocardia concava* strain W9707 (GQ496122.1) | 95 | 95 | 96 | 96 | 96 | 96 | 96 | - |  |  |  |  |  |  |  |  |  |  |  |  |  |  |  |  |
| [9] *Nocardia concava* strain IFM0354 (AB427098.1) | 95 | 95 | 96 | 96 | 96 | 96 | 96 | 100 | - |  |  |  |  |  |  |  |  |  |  |  |  |  |  |  |
| [10] *Nocardia niigatensis* strain IFM0833 (AB450796.1) | 92 | 92 | 92 | 92 | 92 | 92 | 92 | 92 | 92 | - |  |  |  |  |  |  |  |  |  |  |  |  |  |  |
| [11] *Nocardia niigatensis* strain W8186 (GQ496135.1) | 92 | 92 | 92 | 92 | 92 | 92 | 92 | 92 | 92 | 100 | - |  |  |  |  |  |  |  |  |  |  |  |  |  |
| [12] *Nocardia crassostreae* strain IFM10173 (AB450779.1) | 91 | 91 | 91 | 91 | 91 | 91 | 91 | 91 | 91 | 89 | 89 | - |  |  |  |  |  |  |  |  |  |  |  |  |
| [13] *Nocardia otitidiscaviarum* strain NEB252 (CP041695.1) | 90 | 90 | 90 | 90 | 90 | 90 | 90 | 90 | 90 | 89 | 89 | 91 | - |  |  |  |  |  |  |  |  |  |  |  |
| [14] *Nocardia otitidiscaviarum* strain W8030 (GQ496100.1) | 90 | 90 | 90 | 90 | 90 | 90 | 90 | 90 | 90 | 89 | 89 | 91 | 100 | - |  |  |  |  |  |  |  |  |  |  |
| [15] *Nocardia asteroides* strain W7948 (GQ496120.1) | 89 | 89 | 89 | 89 | 89 | 89 | 89 | 88 | 88 | 88 | 88 | 89 | 88 | 88 | - |  |  |  |  |  |  |  |  |  |
| [16] *Nocardia asteroides* strain NCTC11293 (LR134352.1) | 89 | 89 | 89 | 89 | 89 | 89 | 89 | 88 | 88 | 88 | 88 | 89 | 88 | 88 | 100 | - |  |  |  |  |  |  |  |  |
| [17] *Nocardia salmonicida* strain IFM10177 (AB450808.1) | 87 | 87 | 87 | 87 | 87 | 87 | 87 | 87 | 87 | 88 | 88 | 87 | 88 | 87 | 93 | 93 | - |  |  |  |  |  |  |  |
| [18] *Nocardia salmonicida* strain W9876 (GQ496136.1) | 87 | 87 | 87 | 87 | 87 | 87 | 87 | 87 | 87 | 88 | 88 | 87 | 87 | 87 | 92 | 92 | 99 | - |  |  |  |  |  |  |
| [19] *Nocardia mexicana* strain W9754 (GQ496104.1) | 87 | 87 | 87 | 87 | 87 | 87 | 87 | 88 | 88 | 87 | 87 | 87 | 88 | 89 | 89 | 89 | 87 | 87 | - |  |  |  |  |  |
| [20] *Skermania piniformis* strain IFO15059 (AB074927.1) | 83 | 83 | 83 | 83 | 83 | 83 | 83 | 82 | 81 | 82 | 82 | 82 | 83 | 83 | 83 | 83 | 83 | 83 | 83 | - |  |  |  |  |
| [21] *Mycobacterium ulcerans* strain ATCC33728 (AP017624.1) | 80 | 80 | 80 | 80 | 80 | 80 | 80 | 80 | 80 | 79 | 79 | 80 | 81 | 81 | 82 | 82 | 82 | 81 | 80 | 81 | - |  |  |  |
| [22] *Mycobacterium ulcerans* strain SGL03 (LR135168.1) | 80 | 80 | 80 | 80 | 80 | 80 | 80 | 80 | 80 | 79 | 79 | 80 | 81 | 81 | 82 | 82 | 82 | 81 | 80 | 81 | 100 | - |  |  |
| [23] *Mycobacterium marinum* strain ATCC 927 (AP018496.1) | 80 | 80 | 80 | 80 | 80 | 80 | 80 | 80 | 80 | 79 | 79 | 80 | 80 | 80 | 82 | 82 | 82 | 81 | 80 | 79 | 99 | 99 | - |  |
| [24] *Mycobacterium marinum* strain CCUG20998 (CP024190.1) | 80 | 80 | 80 | 80 | 80 | 80 | 80 | 80 | 80 | 79 | 79 | 80 | 80 | 80 | 82 | 82 | 82 | 81 | 80 | 79 | 99 | 99 | 100 | - |

In brackets [ ] are the number used as identifier in the columns.
In parenthesis ( ) are the NCBI accession numbers.

In the phylogenetic tree, the isolates TW29420 and TW295/20 were seen to group within the *Nocardia seriolae* but in a different sub-branch. This subgrouping was well supported with bootstrap value of 90%. The species *Nocardia concava* appeared next to *Nocardia seriolae* as the closest sibling taxa. The *gyrB* phylogenetic tree is presented in Fig. 6

**Figure 6.**
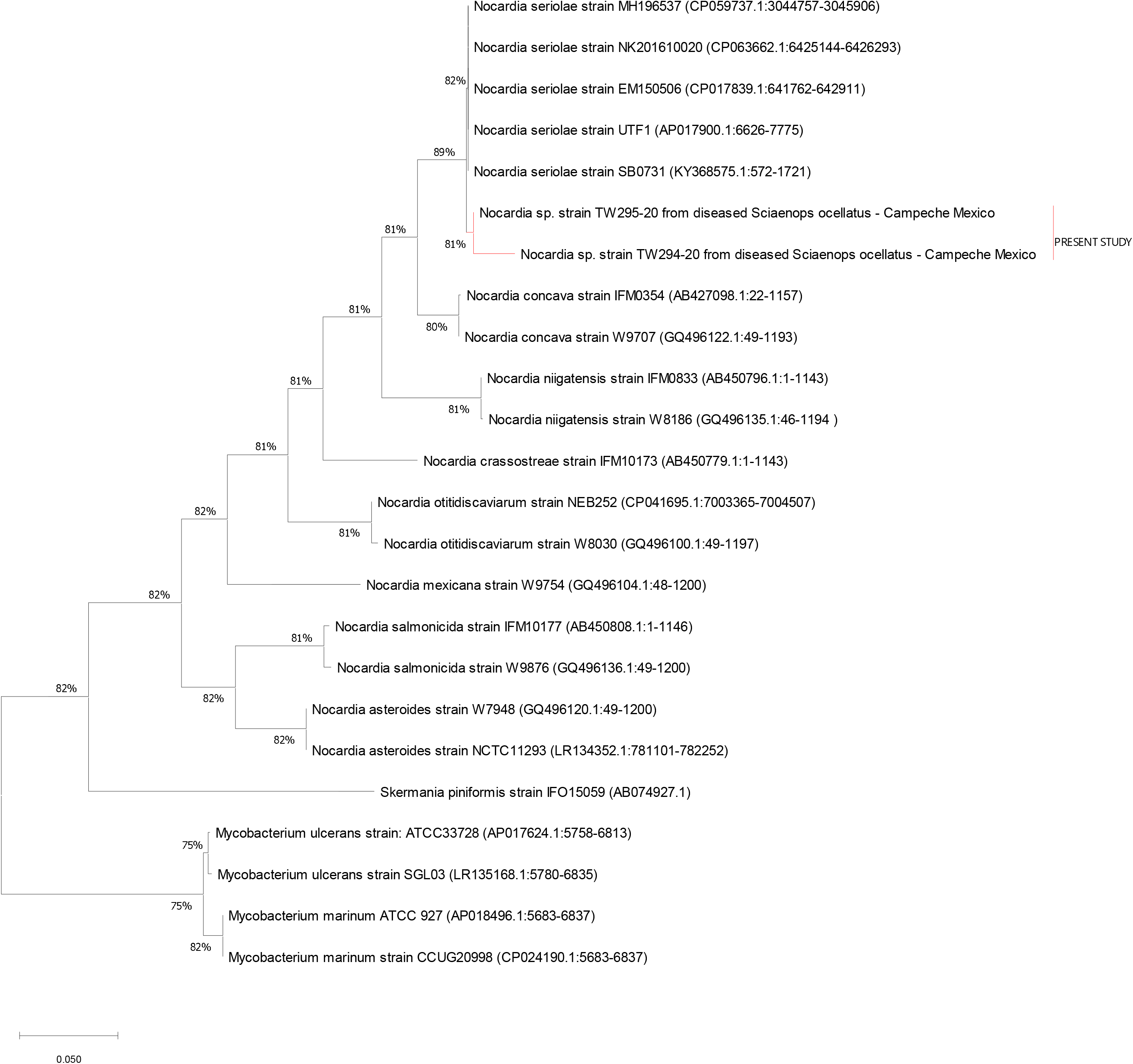
Molecular phylogenetic analysis based on 24 *gyrB* gene sequences (1,411 nt) conducted in MEGA X [Kumar et al., 2018]. The evolutionary history was inferred by using the Maximum Likelihood method and Tamura-Nei model [Kumar and and Nei, 1993]. The tree with the highest log likelihood (−6455.71) is shown. Initial tree(s) for the heuristic search were obtained automatically by applying Neighbour-Join and Bio NJ algorithms to a matrix of pairwise distances estimated using the Tamura-Nei model, and then selecting the topology with superior log likelihood value. The tree is drawn to scale, with branch lengths measured in the number of substitutions per site. The percentage of data coverage for internal node is shown next to the branches.

### Antibiotic resistance profile

Strains CV7Y, CV9Y and CV11Y were resistant to three out of the seven antibiotics tested by the disc diffusion method and displayed medium sensitivity to chloramphenicol and florfenicol, related antibiotics (Table 2). A clear sensitivity to oxytetracycline and doxycycline (derived from tetracycline) was clearly withdrawn from the plate inhibition zones. Doxycycline was used in the grow-out phase during the following years as prophylactic management.

**Table 2.** Antibiotic sensitivity test results of *Nocardia seriolae* by the disc diffusion method.

| Sensi-disc and Concentration | Interpretation |
| --- | --- |
| Doxycycline 30 µg | S |
| Oxytetracycline 30 µg | S |
| Chloramphenicol 30 µg | I |
| Florfenicol 30 µg | I |
| Enrofloxacin (Baytril) 30 µg | R |
| Sulphamethoxazole/Trimethoprim 25 µg | R |
| Clindamycin 2 µg | R |

S=sensitive, I= intermediate, R= resistant

### Disease progression

After the main outbreak in September 2013, a *Nocardia* spp. surveillance programme for the farm was put into practice, involving monthly samples of random collection of fish (n=60) for clinical work, routine bacteriology and testing by PCR. The bacterium was not detected during 2014, likely reflecting the implementation of tighter biosecurity measures and prophylactic applications of medicated feed with Doxycycline. *Nocardia seriolae* colonies were isolated from infected fish displaying mild gross-signs during early and late 2015 only to reappear during mid-2016 and the first semester of 2017. There was no sampling in the second half of 2016 due to company rearrangements priorities. The surveillance sampling was undertaken again in January 2017 but it stopped when farm owners decided to close down business operations. The last two set of samples collected during the spring of 2017 were positive for *N. seriolae*. A summary of the surveillance programme results is presented in Table 3. Overall, there seemed not to be a seasonality effect that favoured outbreaks of nocardiosis in red drum cages, suggesting that there might be other environmental or managerial factors of relevance for the disease persistence.

**Table 3.** Re-emergence of *N. seriolae* in the grow-out phase at the farm between January 2014 and mid-2017.

| Months |  |  |  |  |  |  |  |  |  |  |  |  |
| --- | --- | --- | --- | --- | --- | --- | --- | --- | --- | --- | --- | --- |
| Year | Jan | Feb | March | April | May | June | July | Aug | Sept | Oct | Nov | Dec |
| 2013 |  |  |  |  |  |  |  | - | + | + | + | + |
| 2014 | - | - | - | - | - | - | - | - | - | - | - | - |
| 2015 | + | + | - | - | - | - | - | - | - | + | - | + |
| 2016 | - | - | - | - | - | - | + | No sampling |  |  |  |  |
| 2017 | - | - | - | + | + | End of the surveillance |  |  |  |  |  |  |

+Positive isolation and identification
-No isolation nor detection

## DISCUSSION

In the present study, gross clinical signs along with laboratory and molecular evidence pointed out *Nocardia seriolae* as the etiological agent of the disease. The external lesions as well as the internal gross pathology warranted the inclusion of other white-spot-forming diseases, such as mycobacteriosis, francisellosis, edwardsiellosis and photobacteriosis for the differential diagnosis. The bacteria isolated from the clinically infected fish displayed typical *Mycobacterium* or *Nocardia*-like morphology as observed in tissue imprints, bacterial swabs and histology sections stained with both Gram and Ziehl–Neelsen (ZN) stains.

During the first attempts of identifying the bacterium, PCR primers for *Mycobacteria* spp. were applied giving negative results. Having previously discovered that this was an acid-fast bacterium, these first negative PCR results were the main hint that the outbreak was more likely related to a case of nocardiosis as finally confirmed by our molecular and sequencing results.

*Nocardia* spp. are a Gram-positive, acid-fast, aerobic, non-motile, pleomorphic rod-shaped bacterium that can also be filamentous or branched. Also, researchers have pointed out that some member of the genus *Nocardia* can be partially acid-fast (Muricy et al., 2014), meaning that some parts of the cell wall do not stain, displaying a beaded appearance as presently observed. This characteristic is not commonly present in slow growing *Mycobacterium* spp. Furthermore, histopathology of typical granulomas -some displaying bacterial colonies- and the organs affected, coincides with previous reports of *Nocardia* spp. infections in marine fish (Bransden et al., 2000; Wang et al., 2005; Wang et al., 2009; Cornwell et al., 2011; Elkesh et al., 2013; Wang et al., 2014; Vu-Khac et al., 2016). Most of the above-cited reports describe the liver as one of the commonly affected organs however, in our study the liver, was rarely involved. A very common finding among the diseased specimens though, was liver lipoidosis. We can only speculate that this degrading condition may have acted as a protective factor against the establishment of the bacterium in that particular organ, while liver condition was probably related to a poor nutritional status that in turn could have been the sufficient factor for the persistence of the disease overall. Unfortunately, Koch’s postulates could not be fulfilled due to experimental challenge facilities constraints.

In teleost fish, the *Nocardia* genus contains three pathogenic species of aquaculture importance: *N. seriolae, N. salmonicida* and *N. asteroides*. *N. seriolae* afflicts mostly marine species. It is a well-known fish pathogen in the Easter Asia region including China, Taiwan and Japan (Wang et al., 2005; Labrie et al., 2008; Wang et al., 2014; Vu-Khac et al., 2016). In 1968 was first isolated from the yellowtail *Seriola quinqueradiata* and amberjack *S. dumerili*, (Kariya et al. 1968) but it was identified as *Nocardia kampachi* in posterior discoveries (Ksuda et al. 1974). In 1988 was finally named by Kudo and colleagues (1998) as *N. seriolae* after the fish species was firstly described.

Since 2000, sporadic evidence of its presence in other regions of the world has been reported. For instance, Bransden and co-workers (2000) described a low-prevalence granulomatous infection outbreak during a diet trial in Atlantic salmon (*Salmo salar*) in Australia and isolated a branching bacterium with staining and biochemical characteristics of *N. seriolae*. Ten years later, wild weakfish *Cynoscion regalis* captured by researchers at the University of Ithaca NY (USA) developed chronic mortalities after a month in captivity. *N. seriolae* was identified by comparative sequencing analyses with other *Nocardia* spp. and *Mycobacterium* spp. using DNA obtained from wax-embedded tissues (Cornwell et al., 2011) as the research team did not use LJM to isolate the pathogen, since acid-fast bacterial infection was not suspected. Elkesh et al. (2013) described a systemic outbreak of the disease in cage cultured meagre (*Argyrosomus regius*) from central and western Greece due to *Nocardia* sp. and suggested hatcheries in France and Italy as possible sources. The presence of the pathogen in the later countries is presently unknown.

In the present study, the supply of red drum fry for the farm in Campeche initially had a double source, they were imported from Israel and acquired in a local hatchery. During 2017, the farm developed their own fry production system that remained active until close down. Doxycycline was used as antibiotic in the nursery phase during the cycles following the outbreaks as a prophylactic treatment and showed a significant success in preventing outbreaks due to nocardiosis.

Red drum was introduced into the state of Campeche by the local government since 2006 targeting aquaculture for small holders, but these plans did not prosper although the local hatchery continued active. At the farm of our study, the double-source fry was occasionally mixed during grading, making it difficult to discern if the infection developed locally or the pathogen was imported. As no other outbreak of *N. seriolae* has been reported in Mexico, molecular approaches including whole genome sequencing is required for further comparisons (molecular epidemiology) to elucidate the provenance of the bacteria.

Over 40 marine teleost fish are currently susceptible to nocardiosis by *N. seriolae* infections (Itano et al., 2006a; Elkesh et al., 2013; Sales-Ribeiro, 2016). Labrie et al. (2008) in their review, include an undetermined freshwater tilapia species as susceptible as well. Yet, there seem to be some naturally resistant fish species to nocardiosis; Elkesh et al. (2013) described that the affected meagre, *Aryrosoumus regius* was co-cultured with three other fish species in adjacent net pens, namely seabass, *Dicentrarchus labrax*, seabream, *Sparus aurata* and sharpsnout seabream, *Diplodus puntazzo*, but none of the former two species displayed the lesions observed in meagre. Sharpsnout did display granulomatous lesions in the spleen, but they were attributed to nutritional causes. To their knowledge, the examined fish were not submitted to a recognizable source of stress related to fish farming (e.g. high stocking density, grading or treatment procedures) and speculate that environmental factors may have played a role. The fish farm subject of our study was deployed in what can be considered a pristine environment. Cages were 34 km offshore in crystal-clear waters, 14 m depth, with constant soft currents, no large-vessels traffic and no other fish farms in the surroundings. Furthermore, the northern coasts of Campeche have no large-scale industries or recognizable important pollution discharges and the fishery activity is mostly artisanal. We suspect that the main risk factor was managerial (nutritional deficiency derived of deprived of feeding or inadequate food), that could have been the main sources of stress. Infections can be asymptomatic and, combined with the slow growth rate of *Nocardia*, it can make it difficult to identify the disease in early stages in open aquaculture facilities.

In support of the former hypothesis, Bransden et al. (2000) suggested a combined effect of a poor experimental diet and organic matter built-up in the experimental tanks harbouring *Salmo salar* as the main stress factors for their reported outbreak of *N. seriolae*. In addition, immunological traits among different fish species may play a role for the susceptibility to the disease as it has been suggested that resistant fish species to nocardial infections depend on the production of superoxide anion (respiratory burst) by phagocytes (Kusuda et al., 1989; Itano and co-workers (2006b, c)). Itano and co-workers (2006b, c) performed experimental nocardial infections and found that yellowtail *S. quinqueradiata* was highly susceptible to succumb to the disease as is non-capable of entering in respiratory burst, conversely with the red sea bream *Pagrus major,* a non-susceptible, respiratory-burst capable species.

As a facultative intracellular pathogen, *N. seriolae* can be a rather elusive. Most of the outbreaks reported in the literature reveal that the bacterium was isolated when major gross signs were displayed (Labrie et al., 2008; Wang et al., 2009; Wang et al., 2014); early stages of the infection seem to pass unnoticed, as the invasion of tissue occurs gradually. This is the reason why nocardiosis is not rapidly detected in aquaculture (Wang et al., 2009; Elkesh et al., 2013).

Traditionally, *N. seriolae* has been isolated on brain heart infusion agar (BHIA), tryptone soy agar (TSA) and nutrient agar (NA), with optimum growth at 20–30°C temperature (Kusuda, Taki & Takeuchi, 1974). However, during clinical work, the use of general nutrient media may lead to a wrong diagnosis or miss the true causative pathogen. In our case, *N. seriolae* was first isolated using TSA, as nocardiosis was not suspected and LJ media was not used (Cornwell et al., 2011). In our experience, however, it seems that heavy infection with obvious gross external and internal signs of disease are a condition for TSA and/or LJM successful first isolation, as suggested by Labrie et al. (2008). Since first discovering that an acid-fast bacterium was involved, we included the use of LJM, and during the surveillance (2014-2017), a slightly better rate of isolation was achieved (~45%). Labrie et al. (2008) also experimented on non-recoverable bacteria from some obviously infected fish with a rate of success as low as 4%. They also suggest that there might be a threshold amount of tissue infecting bacteria that allow its recovery through standard methods.

Similarly, Cornwell et al. (2011) did not use LJM during their diagnostic work in a *N. seriolae* outbreak on weakfish and failed to isolate it on marine agar; the pathogen was only revealed by posterior PCR and comparative sequence alignment from infected tissue samples. This highlights the importance of molecular identification and sequencing when dealing with fastidious bacterial pathogens that cause chronic granulomatous disease with heavy cumulative mortality. In the absence of these techniques, laboratories are probably susceptible to fail a proper diagnosis before a nocardial infection in marine fish.

Most of the molecular techniques to identify *N. seriolae* in fish samples are PCRs targeting the 16S rRNA gene (Miyoshi & Suzuki, 2003; Labrie et al., 2008;), although the whole genome sequence of the bacteria is now available (Imajoh et al., 2016; Yasuike et al., 2017: Han et al., 2019). In the present study, the strain J8P8 isolated from diseased red drum was successfully identified as *N. seriolae* using primers to detect a specific region of the 16S rRNA gene (Miyoshi & Suzuki, 2003). However, for a rapid diagnostic test it is necessary to have reliable molecular tools on farm-sites with high sensitivity and sensibility capacities such as the loop-mediated isothermal amplification developed by Itano et al. (2006a).

The genetic characterization of J8P8 strain (isolates TW 294/20 and TW295/20) based on partial *gyrB* gene sequencing allowed to confirm the identity of the pathogen at the species level by gene sequence similarity and to conduct molecular phylogenetic analyses. This, permitted to elucidate the evolutionary relationship of the strain with the closest validly described members in the genus *Nocardia* including an isolate recovered from a human patient in Mexico City (e.g. *Nocardia mexicana*) and other relevant species of the families *Nocardiaceae* and *Mycobacteriaceae*. Interestingly, the closest sibling taxa appeared to be *Nocardia concava* and *Nocardia nigatensis* while the other whole known fish pathogens branched in a distant cluster.

The state of Campeche possesses 521 km of shoreline with a high potential for the development of medium to large-scale marine aquaculture, fully unexplored yet. Most of the current aquaculture endeavours are carried out in in-land freshwater facilities, largely tilapia fish. Mexico possesses laboratories with the standard and advanced tests for the detection of OIE-listed diseases. Nevertheless, the scarce amount of fish pathology professionals, the lack of an advanced and appropriate specific method of identification for such cases as the present, and the difficulty in habilitating personnel in advanced molecular methods in the Gulf of Mexico area, increases the vulnerability of that aquaculture potential. Although most curricula for veterinary studies include aquaculture teaching nowadays, pathology as a subject with sound microbiology knowledge, needs to be reinforced.

Concern on this first isolation of *N. seriolae* and its dispersion arises as the potential of marine aquaculture is near to consolidation in some areas of Mexico (Davila-Camacho et al., 2019) mainly in the states of central and north Pacific coasts. Technology for the culture and growth of several marine species, such as tuna fish *Thunnus orientalis* (Zertuche-Gonzalez et al., 2008), spotted rose snapper *Lutjanus guttatus* (Alvarez-Lajonchere et al., 2012), bullseye puffer *Sphoeroides annulatus* (García-Ortega, 2009), Pacific white snook *Centropomus viridis* (Ibarra-Castro et al., 2017) among others is currently available. Farms with large commercial scale facilities exist at least for two marine teleost fish (the amberjack *Seriola rivoliana* and the sea bass *Morone saxatilis*) in the Peninsula of Baja California (Rueda-López et al., 2017). The establishment of the pathogen subject of the present study in the local fauna and the risk that this may represents, is a matter of further elucidation.

## CONCLUSION

The present study provides a clinical description and thoroughly molecular identification of the first case of nocardiosis in a marine fish aquaculture species in Mexico on the Atlantic side, stringent surveillance must take place in other marine aquaculture development areas of the country. Further research is needed to sequence the whole genomes of *N. seriolae* strains isolated from diseased fish to identify potential virulence and antibiotic resistance genes and conduct molecular epidemiology. In addition, the local dispersion and impact of the pathogen among native fish in the original site of discovery is awaiting to be studied. To the best of our knowledge, no previous reports of nocardiosis have been issued in marine fish aquaculture, wild marine fish species or any other aquatic species in Mexico.

## ACKNOWLEDGEMENTS

The authors would like to thank Ana Delia Cu Escamilla and Karla Aguilar Rendón for their technical support in the lab during the preparation of the histological and species-specific PCR analysis.

## DATA AVAILABILITY STATEMENT

The manuscript does not contain shared data.

## REFERENCES

Alvarez-Lajonchère, L., Abdo De La Parra, M. I., Rodríguez Ibarra, L. E., Velasco Blanco, G., Puello-Cruz, A. C., González Rodríguez, B., & Ibarra Castro, L. (2012). The scale-up of spotted rose snapper, *Lutjanus guttatus*, larval rearing at Mazatlan, Mexico. Journal of the World Aquaculture Society, 43(3), 411–422. https://doi.org/10.1111/j.1749-7345.2012.00573.x

Aubry, A., Mougari, F., Reibel, F., & Cambau, E. (2017). “Mycobacterium marinum.” Tuberculosis and Nontuberculous. Mycobacterial Infections, 735–752. https://doi.org/10.1128/9781555819866.ch43

Aviles-Quevedo A. & Castello-Orvay, F. (2004). An aquaculture manual for *Seriola lalandi* (Pisces: Carangidae) in Baja California Sur, Mexico, National Fisheries Administration of Mexico, 64p (In Spanish). https://www.inapesca.gob.mx/portal/Publicaciones/Manuales/2004-Aviles-y-Castello-Manual-cultivo-Seriola-lalandi.pdf?download

Bransden, M. P., Carson, J., Munday, B. L., Handlinger, J. H., Carter, C. G. & Nowak, B. F. (2000). Nocardiosis in tank-reared Atlantic salmon, *Salmo salar* L. Journal of Fish Diseases 23, 83–85. https://doi.org/10.1046/j.1365-2761.2000.00201.x

Chen, S. C., Lee, J. L., Lai, C. C., Gu, Y. W., Wang, C. T., Chang, H. Y. & Tsai, K. H. (2000). Nocardiosis in sea bass, *Lateolabrax japonicus* in Taiwan, Journal of Fish Diseases 23, 299–307. https://doi.org/10.1046/j.1365-2761.2000.00217.x

Cornwell, E. R., Cinelli, M. J., McIntosh, D. M., Blank, G. S., Wooster, G. A., Groocock, G. H. R., Getchell, G. & Bowser, P. R. (2011). Epizootic Nocardia infection in cultured weakfish *Cynoscion regalis* (Bloch and Schneider), Journal of Fish Diseases 34, 567–571. https://doi.org/10.1111/j.1365-2761.2011.01269.x

Dávila-Camacho, C.A., Galaviz-Villa, I., Lango-Reynoso, F., Castañeda-Chávez, M.D.R., Quiroga-Brahms, C. and Montoya-Mendoza, J., 2019. Cultivation of native fish in Mexico: cases of success. Reviews in Aquaculture, 11(3), pp.816–829. https://doi.org/10.1111/raq.12259

Elkesh, A., Kantham, K. P. L., Shinn, A. P., Crumlish, M., & Richards, R. H. (2013). Systemicnocardiosis in a Mediterranean population of cultured meagre, *Argyrosomus regius* Asso (Perciformes: Sciaenidae). Journal of Fish Diseases, 36(2), 141–149. https://doi.org/10.1111/jfd.12015

FAO (2005-2020). Cultured Aquatic Species Information Programme. Sciaenops ocellatus. Text by Cynthia K. Faulk, A. In: FAO Fisheries Division [online]. Rome. Updated 9 February 2005. [Accessed 27 November 2020]. http://www.fao.org/fishery/collection/cultured-species/en

FAO (2017) National Aquaculture Sector Fact Overview Sheet – México. Text by Montero Rodríguez, M. In: FAO Fisheries and Aquaculture Department [on line], Rome. [Cited April 24, 2017]. http://www.fao.org/fishery/countrysector/naso_mexico/en

Fryer, J.L. & Lannan, C.N., 2015. *Piscirickettsia*. Bergey’s manual of systematics of archaea and bacteria, pp.1–8. https://doi.org/10.1002/9781118960608.gbm01219

García-Ortega, A. (2009). Nutrition and feeding research in the spotted rose snapper (*Lutjanus guttatus*) and bullseye puffer (*Sphoeroides annulatus*), new species for marine aquaculture. Fish Physiology and Biochemistry, 35, 69–80. https://doi.org/10.1007/s10695-008-9226-1

Han, H. J., Kwak, M. J., Ha, S. M., Yang, S. J., Kim, J. D., Cho, K. H., Kim, T. W., Cho, M. Y., Kim, B. Y., Jung, S. H., & Chun, J. (2019). Genomic characterization of *Nocardia seriolae* strains isolated from diseased fish. MicrobiologyOpen, 8(3), e00656. https://doi.org/10.1002/mbo3.656

Hudzicki, J. (2009) Kirby-Bauer disk diffusion susceptibility test protocol. ASM Microbe Library, American Society for Microbiology. https://asm.org/getattachment/2594ce26-bd44-47f6-8287-0657aa9185ad/Kirby-Bauer-Disk-Diffusion-Susceptibility-Test-Protocol-pdf.pdf

Ibarra-Castro, L., Navarro-Flores, J., Sánchez-Téllez, J. L., Martínez-Brown, J. M., Ochoa-Bojórquez, L. A., & Rojo-Cebreros, Á. H. (2017). Hatchery Production of Pacific White Snook at CIAD-Unity Mazatlan, Mexico. World Aquaculture, 25. https://www.researchgate.net/publication/320106950_Hatchery_Production_of_Pacific_White_Snook_at_CIAD-Unity_Mazatlan_Mexico

Imajoh, M., Sukeda, M., Shimizu, M., Yamane, J., Ohnishi, K., & Oshima, S. I. (2016). Draft genome sequence of erythromycin-and oxytetracycline-sensitive *Nocardia seriolae* strain U-1 (NBRC 110359). Genome Announcements, 4(1). https://doi.org/10.1128/genomeA.01606-15

Itano, T., Kawakami, H., Kono, T., & Sakai, M. (2006a). Detection of fish nocardiosis by loop-mediated isothermal amplification. Journal of Applied Microbiology, 100(6), 1381–1387. https://doi.org/10.1111/j.1365-2672.2006.02872.x

Itano, T., Kawakami, H., Kono, T. & Sakai, M. (2006b). Experimental induction of nocardiosis in yellowtail, *Seriola quinqueradiata* Temminck & Schlegel by artificial challenge. Journal of Fish Diseases 29, 529–534. https://doi.org/10.1111/j.1365-2761.2006.00748.x

Itano, T., Kawakami, H., Kono, T. & Sakai, M. (2006c). Live vaccine trials against nocardiosis in yellowtail *Seriola quinqueradiata*. Aquaculture 261, 1175–1180. https://doi.org/10.1016/j.aquaculture.2006.09.006

Itano, T., Nakaoka, N., Kawakami, H., Kono, T. & Sakai, M. (2006d). Comparison of sensitivity between yellowtail *Seriola quinqueradiata* and red sea bream *Pagrus major* to *Nocardia seriolae*. Fish Pathology, 41, 135–139. https://doi.org/10.3147/jsfp.41.135

Kariya, S., Kubota, S., Nakamura, E. & Kira, K. (1968) Nocardiosisin cultured yellowtail and amberjack – I. Bacteriological Investigation, Fish Pathology 3, 16–23 (in Japanese).

Kudo, T., Hatai, K., & Seino, A. (1988). *Nocardia seriolae* sp. nov. causing nocardiosis of cultured fish. International Journal of Systematic and Evolutionary Microbiology, 38(2), 173–178. https://doi.org/10.1099/00207713-38-2-173

Kumar, T. and Nei, M., 1993. Estimation of the number of nucleotide substitutions in the control region of mitochondrial DNA in humans and chimpanzees. Molecular Biology and Evolution, 10(3), pp.512–526. https://doi.org/10.1093/oxfordjournals.molbev.a040023

Kumar, S., Stecher, G., Li, M., Knyaz, C. and Tamura, K., 2018. MEGA X: molecular evolutionary genetics analysis across computing platforms. Molecular Biology and Evolution, 35(6), pp.1547–1549. https://doi.org/10.1093/molbev/msy096

Kusuda, R., Taki, H. &. Takeuchi, T. (1974). Studies on a *Nocardia* infection of cultured yellowtail – II. Characteristics of *Nocardia kampachi* isolated from a gill-tuberculosis of yellowtail, Bulletin of the Japanese Society of Scientific Fisheries 40, 369–373. https://doi.org/10.2331/suisan.40.369

Kusuda, R., Kimura, Y., & Hamaguchi, M. (1989). Changes in peripheral and peritoneal leucocytes in yellowtail *Seriola quinqueradiata* immunized with *Nocardia kampachi*. Bulletin of the Japanese Society of Scientific Fisheries (Japan). https://doi.org/10.2331/suisan.55.1183

Labrie, L., Ng, J., Tan, Z., Komar, C., Ho, E., & Grisez, L. (2008). Nocardial infections in fish: an emerging problem in both freshwater and marine aquaculture systems in Asia. Diseases in Asian aquaculture VI. Fish Health Section, Asian Fisheries Society, Manila, 297–312. http://www.fhs-afs.net/daa_vi_files/22.pdf

Martínez-Lara, P., Martínez-Pochas, M., Gollas Galván, T., Hernández-López, J. & Robles-Pochas G. (2020). Granulomatosis in fish aquaculture: a mini review, Reviews in Aquaculture, 1.20. https://doi.org/10.1111/raq.12472

Miyoshi, Y., & Suzuki, S. (2003). A PCR method to detect *Nocardia seriolae* in fish samples, Fish Pathology, 38 (3): 93–97. https://www.jstage.jst.go.jp/article/jsfp1966/38/3/38_3_93/_pdf

Muricy, E. C. M., Lemes, R. A., Bombarda, S., Ferrazoli, L., & Chimara, E. (2014). Differentiation between *Nocardia* spp. and *Mycobacterium* spp.: critical aspects for bacteriological diagnosis. Revista do Instituto de Medicina Tropical de São Paulo, 56(5), 397–401. https://doi.org/10.1590/S0036-46652014000500005

Nayak, S. K., & Nakanishi, T. (2016). Development of vaccines against nocardiosis in fishes. In Vaccine Design (pp. 193–201). Humana Press, New York, NY. https://doi.org/10.1007/978-1-4939-3389-1_13

Ramirez-Paredes, J. G., Larsson, P., Thompson, K. D., Penman, D. J., Busse, H. J., Ohrman, C., Sjodin, A., Soto, E., Richards, R. H., Adams, A. & Colquhoun, D. J. (2020). Reclassification of *Francisella noatunensis* subsp. *orientalis* Ottem et al. 2009 as *Francisella orientalis* sp. nov., *Francisella noatunensis* subsp. *chilensis* subsp. nov. and emended description of *Francisella noatunensis*. International Journal of Systematic and Evolutionary Microbiology. https://doi.org/10.1099/ijsem.0.004009

Reichley, S.R., Ware, C., Steadman, J., Gaunt, P. S., García, J. C., LaFrentz. B. R., Thachil, A., Waldbieser, G. C., Stine, C. B., Buján, N., Arias, C. R., Loch, T., Welch, T. J., Cipriano, R. C., Greenway, T. E., Khoo, L. H., Wise, D. J., Lawrence, M. L., Griffin, M. J. (2017). Comparative phenotypic and genotypic analysis of *Edwardsiella* isolates from different hosts and geographic origins, with emphasis on isolates formerly classified as *E. tarda*, and evaluation of diagnostic methods. Journal of Clinical Microbiology. 55:3466 –3491. https://doi.org/10.1128/JCM.00970-17

Rueda-López, S., Martínez-Montaño, E. and Viana, M.T., 2017. Biochemical characterization and comparison of pancreatic lipases from the Pacific bluefin tuna, *Thunnus orientalis*; totoaba, *Totoaba macdonaldi*; and striped bass, *Morone saxatilis*. Journal of the World Aquaculture Society, 48(1), pp.156–165. https://doi.org/10.1111/jwas.12372

Sales-Ribeiro, M. C. (2016) Study of the pathogenicity of *Nocardia* spp. by experimental infection in meagre, *Argyrosomus regius*, (Asso, 1801) (in Portuguese), Ph. D. Thesis, Veterinary Faculty, University of Humanities and Technologies, 108p. https://www.semanticscholar.org/paper/Study-of-the-pathogenicity-of-Nocardia-spp.-by-in-Ribeiro/163ecb11c2122ff655c9b2d3793555bc2c80070c

Talaat A.M., Reimschuessel R. & Trucksis M. (1997) Identification of mycobacteria infecting fish to the species level using polymerase chain reaction and restriction enzyme analysis. Veterinary Microbiology 58, 229–237. https://doi.org/10.1016/S0378-1135(97)00120-X

Téllez Castañeda, M. (2019) Perspective of the shrimp aquaculture industry in Mexico II (in Spanish, on-line publication). El Economista. 2019.

Vu-Khac, H., Chen, S. C., Pham, T. H., Nguyen, T. T. G., & Trinh, T. T. H. (2016). Isolation and genetic characterization of *Nocardia seriolae* from snubnose pompano *Trachinotus blochii* in Vietnam. Diseases of Aquatic Organisms, 120(2), 173–177. https://doi.org/10.3354/dao03023

Wang, G. L., Yuan, S. P., & Jin, S. (2005). Nocardiosis in large yellow croaker, *Larimichthys crocea* (Richardson). Journal of Fish Diseases, 28(6), 339–345. https://doi.org/10.1111/j.1365-2761.2005.00637.x

Wang, P. C., Chen, S. D., Tsai, M. A., Weng, Y. J., Chu, S. Y., Chern, R. S., & Chen, S. C. (2009). *Nocardia seriolae* infection in the three striped tigerfish *Terapon jarbua* (Forsskal). Journal of Fish Diseases 32, 301–310. https://doi.org/10.1111/j.1365-2761.2008.00991.x

Wang, P. C., Tsai, M. A., Liang, Y. C., Chan, Y., & Chen, S. C. (2014). *Nocardia seriolae*, a causative agent of systemic granuloma in spotted butterfish, *Scatophagus argus*, Linn, African Journal of Microbiology Research, 8, 3441–3452. https://doi.org/10.5897/AJMR2014.6874

Weisburg, W. G., Barns, S. M., Pelletier, D. A., & Lane, D. J. (1991). 16S ribosomal DNA amplification for phylogenetic study. Journal of Bacteriology, 173(2), 697–703. https://jb.asm.org/content/173/2/697.short

Woo, P. T. K. & Bruno, D. W. (2011). Fish Diseases and Disorders III: Viral, Bacterial and Fungal Infections, CAB Publishing, 940 p. https://www.cabi.org/bookshop/book/9781845937379/

Yamamoto, S., & Harayama, S. (1995). PCR amplification and direct sequencing of *gyrB* genes with universal primers and their application to the detection and taxonomic analysis of *Pseudomonas putida* strains. Applied and Environmental Microbiology, 61(3), 1104–1109. https://www.ncbi.nlm.nih.gov/pmc/articles/PMC167365/

Yasuike M, Nishiki I, & Iwasaki Y. (2017). Analysis of the complete genome sequence of *Nocardia seriolae* UTF1, the causative agent of fish nocardiosis: The first reference genome sequence of the fish pathogenic *Nocardia* species. PLoS One. 12(3):e0173198. https://doi.org/10.1371/journal.pone.0173198

Zertuche-González, J. A.; Sosa-Nishizaki, O.; Vaca Rodriguez, J. G.; del Moral Simanek, R.; Yarish, C., & CostaPierce, B. A. (2008). Marine Science assessment of capture-based tuna (*Thunnus orientalis*) aquaculture in the Ensenada region of northern Baja California, Mexico. Publications.https://opencommons.uconn.edu/cgi/viewcontent.cgi?article=1000&context=ecostam_pubs

